# Context-dependent effects of temperature and community complexity on trophic interaction strength in a mite community

**DOI:** 10.1101/2022.12.22.521664

**Authors:** Inmaculada Torres-Campos, Sara Magalhães, Jordi Moya-Laraño, Marta Montserrat

**Affiliations:** Instituto de Hortofruticultura Subtropical y Mediterránea “La Mayora”, Universidad de Málaga, Consejo Superior de Investigaciones Científicas (IHSM-UMA-CSIC), Avda Dr. Weinberg s/n, Algarrobo-Costa, 29750 Málaga, Spain; cE3c: Centre for Ecology, Evolution and Environmental Changes, Faculdade de Ciências, Universidade de Lisboa, Campo Grande, Edifício C2, 1749-016 Lisboa, Portugal; Estación Experimental de Zonas Áridas - CSIC, Carretera de Sacramento s/n, La Cañada de San Urbano, 04120 Almería, Spain

## Abstract

Both temperature and community complexity are known to affect trophic interaction strength (TIS), but whether their effect is additive or not is as yet an open question. Here we used a 2-predator/3-prey system consistently occurring in avocado orchards to explore the effects of increasing warming and community complexity on the strength of predator:prey trophic interactions. The two predator species differed in their diet breath (a *carnivore* and an *omnivore*) and the prey types included a herbivore, heterospecific juvenile predators, and/or pollen. Overall, analyses revealed multiplicative effects of community complexity and both temperature and predator diet breath on the net predator:prey(s) TIS. Indeed, warming led to increased TIS in the community with omnivore as top predator, but only in absence of its preferred food source. When the carnivore was the top predator, in contrast, higher temperatures led to TIS being more negative, but only for the IGprey. We conclude that sources of context dependence in specific systems need to be identified to unveil effects of warming on communities.

## Introduction

Trophic interactions (e.g. predation, herbivory) are central drivers of the abundance, dynamics and persistence of populations in nature, and play a pivotal role in shaping the structure of ecological communities (Abdala-Roberts et al. 2019). They are the key component of topological food webs ((Paine 1980), Smith and Smith 2009), which describe the trophic links among species in a community and the mechanistic pillars upon which relevant indirect interactions operate, such as those leading to trophic cascades (Schmitz, Hamback, & Beckerman, 2000) or apparent competition (Holt, 1977). Trophic interactions are also used in food web theory to define the degree of complexity in communities (May 1974, Pimm et al 1991). For example, connectivity, connectance, and linkage density are measures of community complexity that consider either the total number of possible trophic interactions, the proportion of realized interactions among all possible ones, or the average number of trophic links per species, respectively (Landi et al, 2018). Hereon, we will refer to community complexity as communities in which complexity is measured via their connectivity, i.e. via the number of potential trophic links.

Theoretical studies based on empirical data show that trophic interaction strength (TIS), here defined as ‘the likelihood of consumption of one species by another’ (McCann, Hastings & Huxel 1998; Berlow et al. 2004), is regulated by community complexity and temperature. On the one hand, increasing the number of species in a community may lead to an increase in the number of potential trophic links among them, which may weaken the overall strength of pair-wise interactions. Indeed, several works state that communities are typically composed of few strong interactions embedded in a preponderance of weak links (O’Gorman and Emmerson 2009, Neutel et al. 2002, Montoya and Sole 2003), and that weak interactions are key to stabilize complex food webs (Paine 1992; McCann et al. 1998; Neutel, Heesterbeek & de Ruiter 2002; Neutel & Thorne 2014, Emmerson et al 2004, Gellner and McCaan 2012, 2016). On the other hand, the negative correlation between the number of potential trophic links among species and the strength of trophic interactions can be inverted when higher-order interactions; that is, interactions in which one species modulates the interaction among other species, are included into models (Bairey et al 2016, Gibbs et al 2022).

In communities composed of ectothermic organisms (e.g. arthropods), moderately high temperature is a key determinant of TIS in both predator-prey and herbivore-plant interactions because it affects metabolic demands of individuals (Brown et al. 2004; Hoekman 2010, Schmalhofer 2011 Schulte 2015) and predator and prey mobility (Brown et al. 2004; Moya-Laraño 2010; Moya-Laraño et al. 2012, Sentis et al 2014). Furthermore, exposure to sufficiently high temperatures; i.e., beyond optimal, may denaturalize proteins and cause death of individuals (Hazell et al 2010) or affect severely their fitness (Roux et al 2010) and performance (Dell et al 2011), thereby reducing TIS.

The effects of temperature and community complexity on TIS is, in general, expected to correlate negatively because higher trophic levels are often more sensitive to warming than lower trophic levels (Petchey et al 1999, Cagnolo et al 2002, Voight et al 2003, Preisser and Strong 2004, Urban et al 2017). However, studies also indicate that the interaction between the two factors is likely to be context dependent. For example, in simple herbivore-plant bi-trophic systems warming can affect negatively primary producers because it tends to strengthen herbivory by accelerating herbivore metabolic rates, thereby accelerating their feeding, growth and reproduction rates (O’Connor et al. 2009, Zidon et al 2016). However, plants are also ectotherms, and in some circumstances (e.g., high water availability) their biomass could grow faster than that of their consumers at higher temperatures (Brown et al. 2004).

In linear tri-trophic systems there are examples in which warming may strengthen top-down trophic cascades because top predators increase their trophic activity (e.g. Katrina et al 2012 in aquatic systems, Barton et al 2009, Frank and Brambock, 2016, El-Danasoury and Iglesias-Piñeiro 2018 in terrestrial systems), or because it leads to a lower proportion of herbivore migrant phenotypes, thereby strengthening predator TIS locally (Wang et al. 2017). Other studies, however, find that warming reduces the strength of trophic interactions, for example because it negatively affects the performance of predators (Dell et al 2011) or their competitive abilities (Guzmán et al 2016).

In other complex community configurations, the effects of temperature on TIS are also variable (Tylianakis et al 2008, Barton et al. 2009, Barton and Schmitz 2009, Harley 2011). For example, in an aquatic system with two predators differing in their foraging strategy (active vs. sit-and-wait) and sharing a prey, warming increased the swimming speed of the prey, which led to a decrease in the TIS in the active predator, but an increase in the TIS in the sit-and-wait predator (Twardochlew et al 2020). Sentis et al (2014) combined experiments and modelling to test the effects of warming and enrichment in a terrestrial tri-trophic food web with intraguild (IG hereafter) predation. They found that enrichment decreased species interaction strength and omnivory, whereas warming strengthened TIS between the IGpredator and both the IGprey and the herbivore. Similarly, Volker & Roman (2018) tested the effects of warming on three communities of increasing complexity using a keystone predator community configuration composed of two species of herbivore frogs sharing a predator and competing for two species of primary food sources. They found that the effect of warming on TIS and non-TIS (competition), and subsequent cascading effects on primary resources, varied across community complexity and strongly depended on species identity.

Here we focus on two potential sources of context dependence inherent in the characteristics of the species of our experimental system (see below), i.e. diet breath of predators and the presence of preferred food sources for predators, which could modulate the relationship between temperature and community complexity on TIS. For example, it is often assumed that species with broad diet breath will have higher chances to adapt to environmental change than species with limited diet breath because the former can incorporate a wider range of potentially available food sources into their diet (Symondson et al. 2002). Yet, it also known that consumers usually seek and exploit first their preferred food sources (Eubanks and Denno 2000). Thus, the presence of preferred food sources for the members of a community could strengthen TIS via the reduction in the number of realized predator-prey links (Torres-Campos et al 2020).

In this work, we used a two predator / three food-source system that naturally occurs in avocado orchards of Southeastern Spain to explore the effects of warming and community complexity, the later defined by the number of potential trophic links in each community (connectivity), on the strength of trophic interactions between predators and their prey(s). In our experimental system, the two species of predators can effectively forage on herbivore prey and on each other’s juveniles (Torres-Campos et al 2020), thus they can potentially engage in reciprocal intraguild predation (Polis et al 1989). Furthermore, the two predators differ in their diet breath and food preferences, with one species (*Neoseiulus californicus*, Acari: Phytoseiidae) being more of a *carnivore* that prefers herbivore prey over heterospecific predator juveniles (i.e. IGprey), but that it is able to use a food source of plant origin, pollen, to survive when animal prey is not available (see experimental system, below); and the other species (*Euseius stipulatus*, Acari: Phytoseiidae) being more of an *omnivore* that exploits both pollen and animal prey, but that prefers the former over the latter. Thus, our experimental system included three potential food sources for the two predators, each of them being a preferred food source for either predator species. By measuring the predation rates of predators on each type of prey, and the concomitant rates of food conversion into eggs, when predators were allowed to forage on an increasing number of food types and across an increasing range of temperatures, we were able to assess (a) whether the overall TIS on the herbivore or on the IGprey changes with temperature, community complexity, and their potential interaction, and (b) whether the impact of these variables on TIS hinges upon the diet breadth of predators or the presence of the preferred food source for each of the predators.

## Material and Methods

### Experimental system

We used a 4-species arthropod community that typically occurs in avocado orchards of South-Eastern Spain. This community is composed of two predator mite species (*Euseius stipulatus* and *Neoseiulus californicus*, Acari: Phytoseiidae), one herbivore mite species (*Olygonychus perseae*, Acari: Tetranychidae), and alternative food in the form of anemophilous pollen of several species deposited in the surface of avocado leaves (González-Fernández et al 2009).

*Euseius stipulatus*, the *omnivore* species, is a generalist predator that forages on soft-bodied animal prey and it contributes to the natural control of tetranychid mites in different fruit crops, such as citrus (Aguilar-Fenollosa et al. 2011a, b; Pascual Ruiz et al. 2014; Pérez-Sayas et al. 2015) and avocado (González-Fernández et al 2009, Maoz 2011, Montserrat et al 2013). However, this species forages preferentially on pollen when it is available as it is its optimal food source (Bouras and Papadoulis 2005; Ferragut et al.1987). *Neoseiulus californicus*, the *carnivore* species, is a specialist predator of tetranychid mites, contributing to the natural control of several spider-mite pest species in several crops (Montserrat et al 2008, Aguilar-Fenollosa et al. 2011). Yet, this species can utilize pollen as food source when animal prey is scarce, or not available, to survive (Pascua et al 2020). Both predator species can potentially engage in size-dependent intraguild predation, with larger individuals (usually females) acting as IGpredators, and heterospecific juvenile stages acting as IGprey (Torres-Campos et al 2020, Urbaneja-Bernat et al. 2019)). *Olygonichus perseae* is a herbivore specific to avocado plants. It was first detected in avocado orchards of Spain in 2006, and it has successfully established and become a pest ever since (Vela et al 2007).

### Mite cultures

Cultures of the three mite species were kept in climate chambers at 25±1°C, 65±5% RH and 16:8h L:D (Light:Dark). Cultures of the *omnivore* mite *E. stipulatus* were started in 2007 from ca. 300 individuals collected from avocado trees located in the experimental station of “La Mayora”. Rearing units consisted of three bean plants (*Phaseolus vulgaris* L.) with 6-10 leaves, positioned vertically, with the stems in contact with sponges (*ca*. 30 x 20 x 5 cm) covered with cotton wool and a plastic sheet (27 x 17 cm), and placed inside water-containing trays (8 L, 42.5 x 26 x 7.5 cm). The plant roots were in contact with the water, and the aerial parts were touching each other, forming a tent-like three-dimensional structure, where individuals could easily walk from one plant to the other. Cotton threads were placed on the leaves, to serve as oviposition sites for the females. Mites were fed *ad libitum* twice a week with pollen of *Carpobrotus edulis* (cat’s claw) spread on leaves with a fine brush. *Euseius stipulatus* is able to develop and reproduce on this food source (Ferragut *et al*. 1987). Every three weeks, new rearings were made by transferring leaves with mites and the cotton threads filled with eggs to a new unit.

The initial population of the *carnivore* mite *N. californicus* was obtained from Koppert Biological Systems S.L. in bottles of 1000 individuals (Spical^®^). Colonies were kept on detached bean leaves infested with *Tetranychus urticae* that were placed on top of inverted flower pots (20 cm Ø) inside water-containing trays, in a climate room at 25+/- 5°C.

The herbivore *Oligonychus perseae* was not maintained in a laboratory culture due to technical difficulties in preserving detached avocado leaves. Instead, they were collected from the field on a regular basis from avocado orchards located in the experimental station of “La Mayora”.

Pollen of *C. edulis* was obtained from flowers collected in the experimental station. Stamens dried in a stove at 37°C for 48h, then sieved (350 μm).

### Experimental design

The experiment was designed to evaluate whether the strength of trophic interactions changes with increasing (1) temperature and (2) community complexity, and whether sources of context-dependency such as (3) diet breadth of predators and (4) presence of preferred food sources for either the predators or the IG prey, contribute to explain the changes. We did this by building community modules that differed in predator and prey identity, and exposed them to different abiotic/biotic treatments. Specifically, (i) temperature varied across three levels, in which individuals experienced either: a) 25°C during the day (16h) and 22°C during the night (8h) (average/h temperature per day: 24°C; hereafter ‘mild’); (b) 30°C during the day and 27°C during the night (average/h temperature per day: 29°C; hereafter ‘hot’); c) or 33°C during the day and 30°C during the night (average/h temperature per day: 32°C; hereafter ‘very hot’). The ‘hot’ condition was obtained averaging daily day/night temperatures registered in July and August over five consecutive years (2006-2010) in the study area (Montserrat et al 2008). The other conditions were modified to mimic local daily/night temperatures typical in spring (‘hot’-5°C=‘mild’) and climate change scenarios (‘hot’+3°C = ‘very hot’) (Torres-Campos et al 2017); (ii) Community complexity was not defined by the number of species but by the number of potential predator-prey trophic links typically depicted in four community modules (sensu Holt 1997, see experimental design in Figure 1) of increasing complexity. We created community modules with 1, 2, 3, or 4 trophic links using different combinations of species and developmental stages: communities with 1 trophic link contained the top predator and either the herbivore or the IGprey; communities with 2 trophic links contained the top predator, pollen, and either the herbivore or the IGprey; communities with 3 trophic links contained the top predator, the IGprey, and the herbivore; and communities with 4 trophic links contained the top predator, the IGprey, the herbivore, and pollen. (iii) Diet breadth was determined by which of the two species acted as the top predator, that is, females of the *omnivore* predator *E. stipulatus* or females of the *carnivore* predator *N. Californicus*. Importantly, (iv) each community included or not a preferred food source for either species, that is, pollen for females and juveniles of the *omnivore* predator, and herbivore prey for females and juveniles of the *carnivore* predator. Throughout the text, the identity of (IG)predators will be indicated as *omnivore* or *carnivore*, and that of IGprey will be indicated as *omnivore*-IGprey or *carnivore-IGprey*.

**Figure 1.**
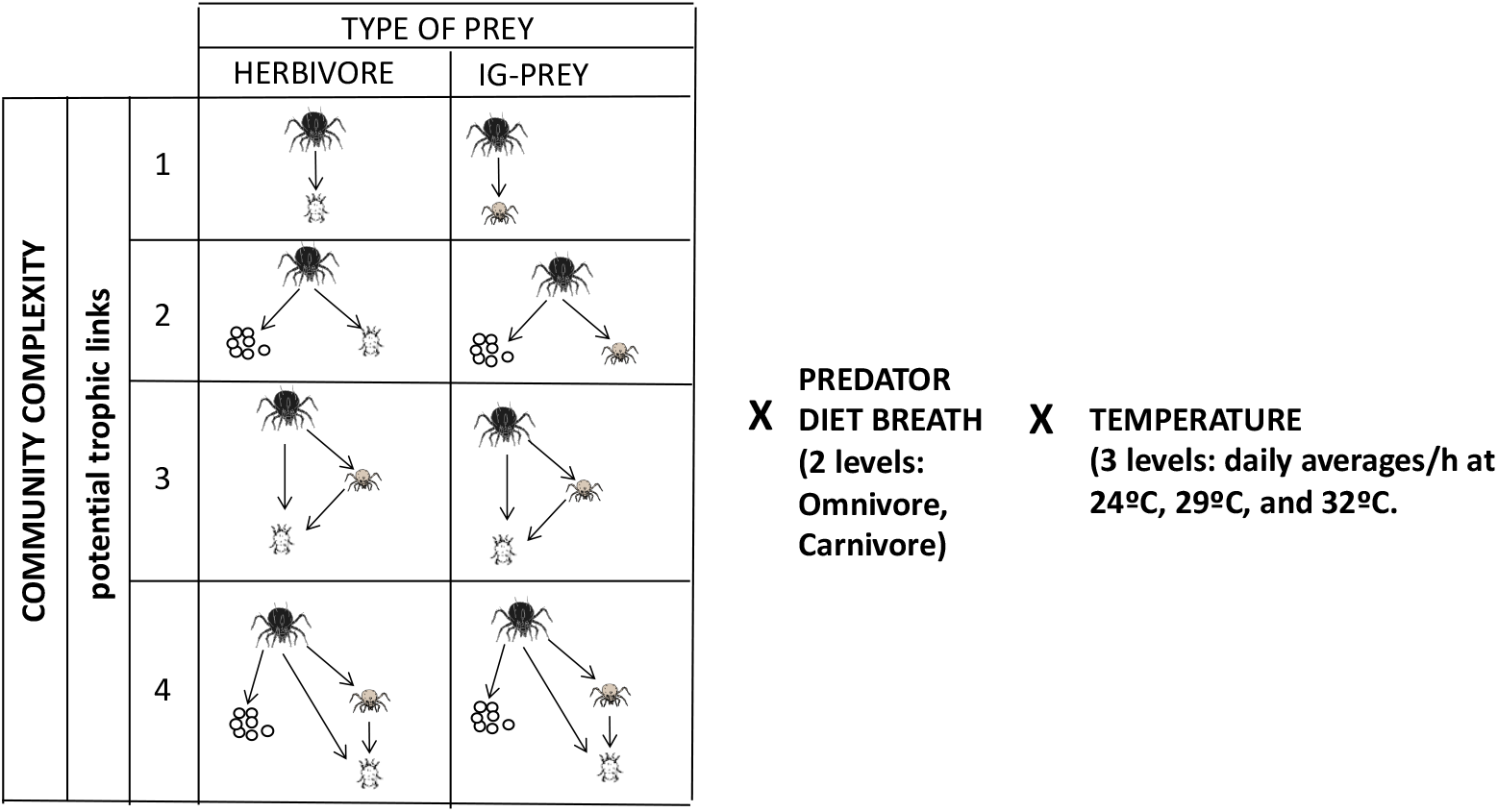
Experimental design: We used an experimental system composed of two species of predatory mites that can potentially engage in reciprocal IGP: *E. stipulatus*, an *omnivore* species that exploits and forages on both soft body arthropods and on sources of vegetal origin (pollen) and *N. californicus*, a *carnivore* species that forages mostly on animal food sources. The experiment was designed to measure trophic interaction strength (TIS) on either the herbivore or the IGprey, in communities of increasing complexity and across different temperatures

### Experimental arenas

Arenas where predators, (IG)prey, and pollen were allowed to interact x have been described in detail in Guzmán *et al*. (2016a). Briefly, a hole (6.5 cm Ø) was cut in a petri dish (9 cm Ø), turned upside down, and filled with an avocado leaf disc (7.5 cm Ø). The borders were glued to a clay ring. Inside the petri dish, wet cotton wool ensured enough humidity to keep leaves turgid. Petri dishes were then sealed with parafilm^®^. To prevent individuals from escaping, a ring of Tanglefoot^®^ was applied along the outer margin of the leaf disc. These arenas were designed to create two abiotic environments: the upper environment, where interacting individuals experience the conditions of treatments, and the lower environment, which is kept humid to maintain the avocado leaf-discs turgor.

Each arena contained one gravid female (10-14 days after egg hatching) as the top predator, which was previously starved for 16 h to standardize hunger among individuals. Arenas allocated to treatments with IG-prey contained 10 predator juveniles (2-3 days old since hatching). Arenas allocated to treatments with the herbivore were built as follows: ten females of *O. perseae* were let to build nests and lay eggs on the arenas for four days. Arenas allocated to treatments with alternative food contained pollen of *C. edulis* supplied *ad libitum* using a fine brush. 24 h after introducing all species, the number of dead herbivores/IG-prey, and the number of eggs laid by female predators, were recorded. Arenas with the herbivore or the IG-prey with neither food nor predators were prepared as control for non-trophic mortality, to obtain an average of natural mortality for either prey at each temperature. This was needed to calculate trophic interaction strengths (see below). Each treatment was replicated between 10 and 18 times.

### Data analyses

The metric for trophic interaction strength (TIS) used was the log response ratio (Berlow et al 2004), defined as ln(# *prey_i,Tj_/# prey_f,Tj_*), where *prey_i,Tj_* is the average number of prey (either the herbivore or the IG-prey) alive after 24 h when trophic mortality was not allowed, and thus natural mortality in the absence of food and predators is considered, at each temperature *Tj;* and *prey_f,Tj_* is the number of prey (either the herbivore or the IG-prey) alive after 24 h interacting with the (IG)predator, at each temperature *Tj*. Thus, all values accounted for natural mortality of (IG)prey at each temperature. This way, TIS > 0 indicates that predation on either the IG-prey or the herbivore prey is the main cause of the reduced number of survivors at the end of the experiment; TIS = 0 indicates that no significant predation occurred; and TIS < 0 indicates that survival of prey, either the IGprey or the herbivore, is higher than the control treatment with no predator or alternative food, probably because of a combination of not being preyed upon and of being able to forage on other available food sources. This metric does not depend on equilibrium conditions and thus it works well for short-term experiments such as ours (Berlow et al 2004). We obtained measures of TIS on either the herbivore or the IGprey exerted by either the *omnivore* or the *carnivore* predator, across an increasing number of potential trophic links in the community, and across an increasing range of temperatures.

Trophic interaction strength (TIS) and oviposition rates of female predators were analysed separately using Generalized Lineal Models (GLM) with a Gaussian/Poisson distribution of errors and identity/log link functions, respectively. For all the applied models we followed a backward elimination procedure: if the higher order interaction among the explanatory variables was not significant and the model without the interaction had lower AIC by more than 2 units, this interaction was removed from the model. Subsequently, the same procedure was sequentially followed for lower order interactions (Crawley 2014). GLM analyses were performed using the “car” package in R and P values were obtained based on type III Likelihood Ratio tests (R Core Team 2017). Post-hoc tests were done, when needed, using the library “multcomp” and Tukey tests.

To explore (a) whether the effects of temperature and community complexity on the TIS experienced by either type of prey, and the concomitant effects on the predator oviposition rates, depended on the diet breadth of the predator, we subjected data (TIS, oviposition rates) to 3-factor GLMs with “temperature” as a continuous variable and “community complexity” and “diet breath” of predators as categorical variables. To explore (b) whether TIS experienced by herbivores confronted to the *omnivore* predator depended on temperature, presence of *carnivore*-IGprey, and/or presence of the preferred food source for the *omnivore* (i.e. pollen), we subjected data from the omnivore species only to 3-factor GLM with “temperature” as continuous variable, and “presence of IGprey” (yes, no) and “presence of preferred food for the predator” f(yes, no) as categorical variables. To explore (c) whether TIS experienced by herbivores confronted to the *carnivore* predator depended on temperature, presence of IG prey, and/or presence of the preferred food source for the IGprey (i.e pollen), we subjected data from the carnivore species only to 3-factor GLM with “temperature” as continuous variable, and “presence of IGprey” and “presence of preferred food for the IGprey” as categorical variables. Last, to explore (d) whether TIS experienced by the IGprey when confronted to either the *carnivore* or the *omnivore* depended on temperature, and on the presence of the preferred food for either the predator or the IGprey, we subjected data to GLMs containing “temperature” as continuous variable, and “presence of preferred food for the predator” and “presence of preferred food for the IG prey” (yes, no) as categorical variables, for each predator species separately.

## Results

### Effects of temperature, community complexity, and diet breadth of predators on overall TIS

Overall TIS on the herbivore was affected by multiplicative effects between temperature and community complexity, and between diet breadth and community complexity (Table 1a). The first interaction was the result of temperature affecting positively TIS on the herbivore, but only in the community with three trophic links (community 3 in figs. 2 and 4, left panels). The second interaction was the result of community complexity affecting TIS on the herbivore, but only when the predator was the *omnivore* (compare left panels in figs. 2 and 4).

**Figure 2.**
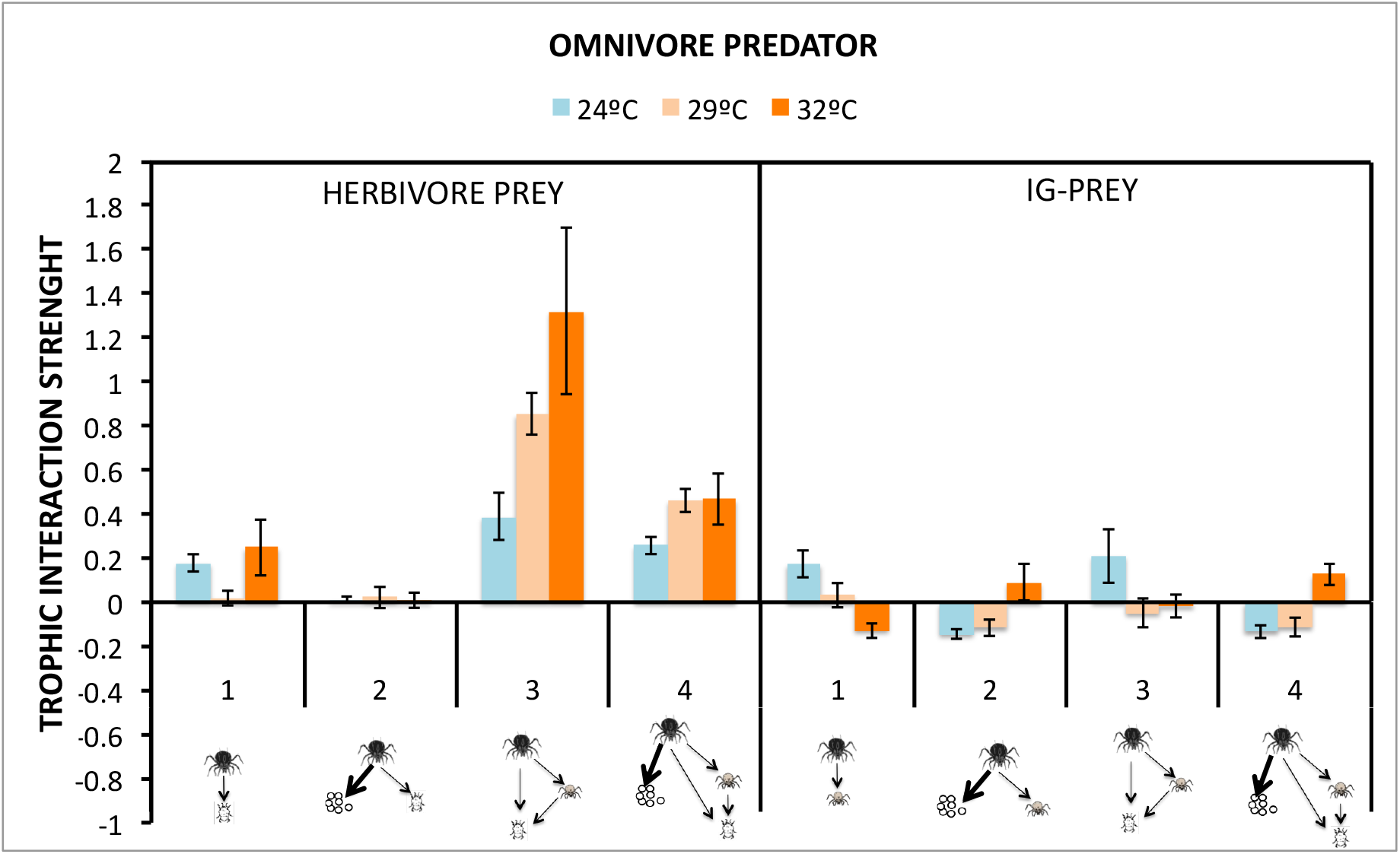
Trophic interaction strength (TIS) exerted by the *omnivore* predator on either the herbivore (left panel) or the *carnivore-IGprey* (right panel), measured in communities of increasing complexity (1 to 4 potential trophic links) and across increasing temperatures.

**Figure 3:**
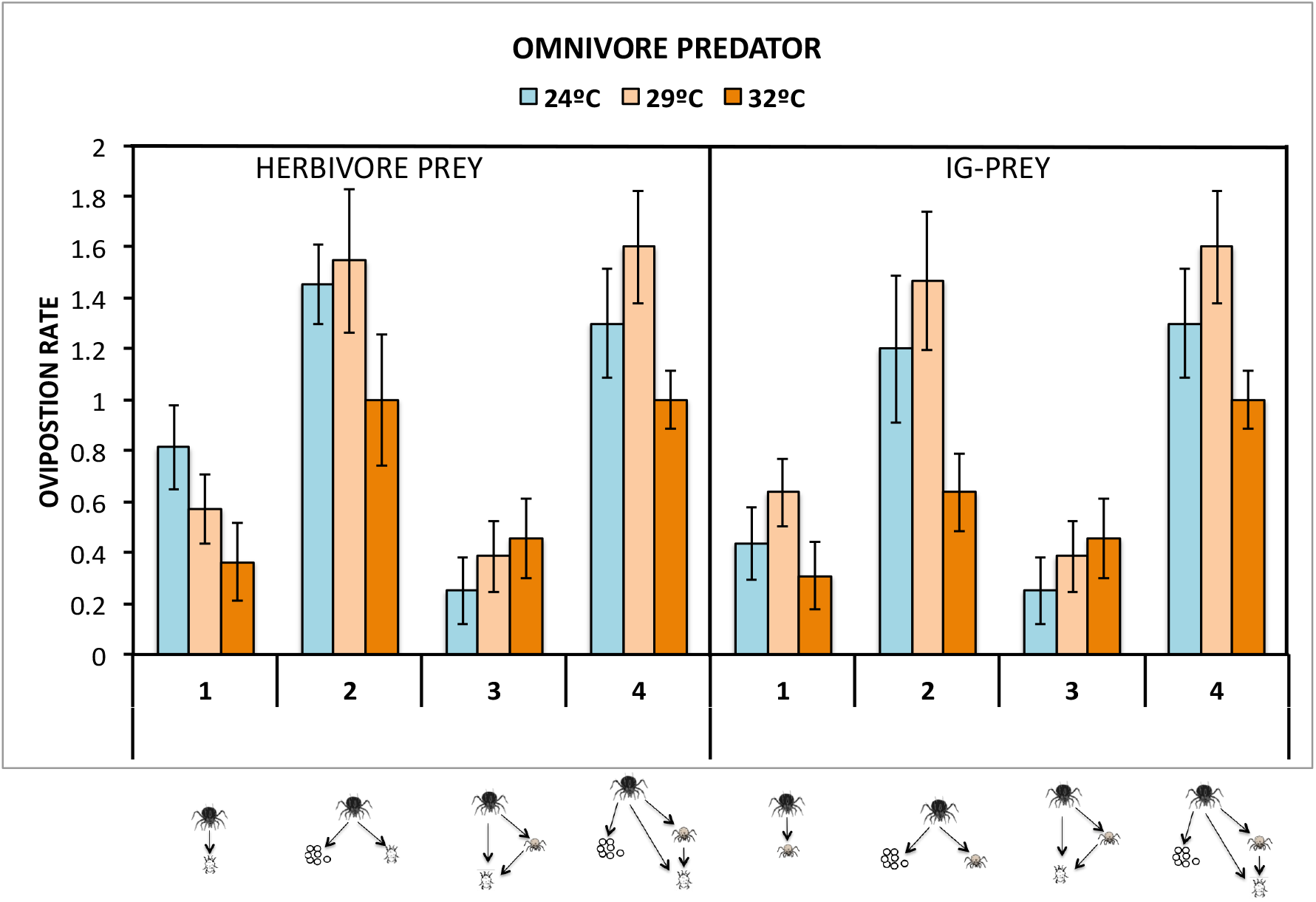
Oviposition rates of the *omnivore* predator when the focal prey was either the herbivore (left panel) or the *carnivore-IGprey*, measured in communities of increasing complexity (1 to 4 potential trophic links) and across increasing temperatures.

**Figure 4.**
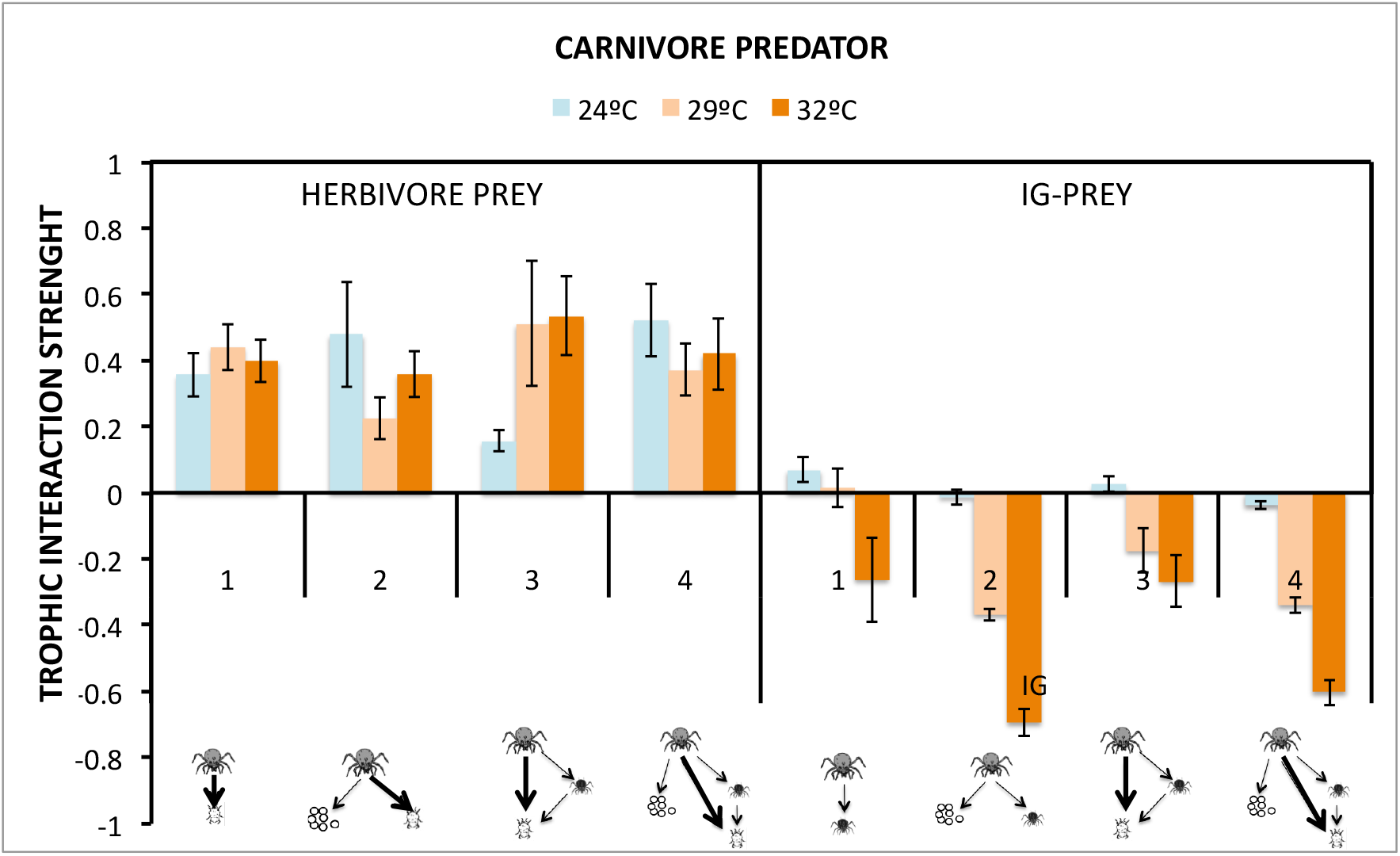
Trophic interaction strength (TIS) exerted by the *carnivore* predator on either the herbivore (left panel) or the *omnivore*-IGprey (right panel), measured in communities of increasing complexity (1 to 4 potential trophic links) and across increasing temperatures.

**Table 1.** Results of the overall Generalized Linear Models applied to the dependent variables “trophic interaction strength” (TIS, with gaussian distribution of errors), and predator’s “oviposition rates” (with poisson distribution of errors), when the prey was either the herbivore (Table 1a and 1b, respectively) or the IG-prey (Table 1c and 1d, respectively). Models included Temperature (3 levels) as continuous explanatory variable, and Community complexity (4 levels) and “Diet Breath” of predators [Omnivore: *Euseius stipulatus*, Carnivore: (*Neoseiulus caiifornicus)]* as categorical variables.

|  | LR Chisq | Df | Pr(>Chisq) |
| --- | --- | --- | --- |
| Temp | 0.189 | 1 | 0.66357 |
| Diet_Breath | 5.796 | 1 | 0.01606 * |
| Com_complex | 22.609 | 3 | 4.871e-05 *** |
| Temp:Diet_Breath | 2.856 | 1 | 0.09104 . |
| Temp:Com_complex | 22.741 | 3 | 4.574e-05 *** |
| Diet_Breath:Com_complex | 46.074 | 3 | 5.469e-10 *** |
| Temp*Diet_Breath*Com_complex | EXCLUDED |  |  |

Table 1b) Overall Oviposition rates (herbivore prey)
|  | LR Chisq | Df | Pr(>Chisq) |
| --- | --- | --- | --- |
| Temp | 2.125 | 1 | 0.14490 |
| Diet_Breath | 5.220 | 1 | 0.02233 * |
| Com_complex | 54.066 | 3 | 1.086e-11 *** |
| Temp*Diet_Breath | EXCLUDED |  |  |
| Temp*Com_complex | EXCLUDED |  |  |
| Diet_Breath*Com_complex | EXCLUDED |  |  |
| Temp*Diet_Breath*Com_complex | EXCLUDED |  |  |

Table 1c) Overall TIS on IG prey
|  | LR Chisq | Df | Pr(>Chisq) |
| --- | --- | --- | --- |
| Temp | 13.168 | 1 | 0.0002848 *** |
| Diet_Breath | 0.051 | 1 | 0.8212065 |
| Com_complex | 10.288 | 3 | 0.0162716 * |
| Temp:Diet_Breath | 0.000 | 1 | 0.9894696 |
| Temp:Com_complex | 16.311 | 3 | 0.0009789 *** |
| Diet_Breath:Com_complex | 46.175 | 3 | 5.206e-10 *** |
| Temp:Diet_Breath:Com_complex | 52.015 | 3 | 2.973e-11 *** |

Table 1d) Overall oviposition rates (IG prey)
|  | LR Chisq | Df | Pr(>Chisq) |
| --- | --- | --- | --- |
| Temp | 0.071 | 1 | 0.790607 |
| Diet_Breath | 7.648 | 1 | 0.005685 ** |
| Com_complex | 5.478 | 3 | 0.139971 |
| Temp:Diet_Breath | 0.560 | 1 | 0.454113 |
| Temp:Com_complex | 1.131 | 3 | 0.769694 |
| Diet_Breath:Com_complex | 32.858 | 3 | 3.45e-07 *** |
| Temp*Diet_Breath*Com_complex | EXCLUDED |  |  |

Trophic interaction strength on the <u>IGprey</u> was influenced by multiplicative effects from all three main factors (Table 1c). This effect was caused by a low effect of temperature and community complexity on the *carnivore*-IGprey’s TIS when the predator was the *omnivore* (figs 2, right panel), and by a positive effect on the survival of the *omnivore*-IGprey as temperature and community complexity increased when the predator was the *carnivore* (fig. 4, right panel). Note that TIS in right panel of figure 4 go from zero to increasing negative values. Oviposition rates, however, depended on the identity of the predator and on community complexity (Table 1d), because overall oviposition rates were higher for the *omnivore* than for the *carnivore*, and differed between the two predator types depending on the community complexity (compare right panels in figs 3 and 5).

**Figure 5:**
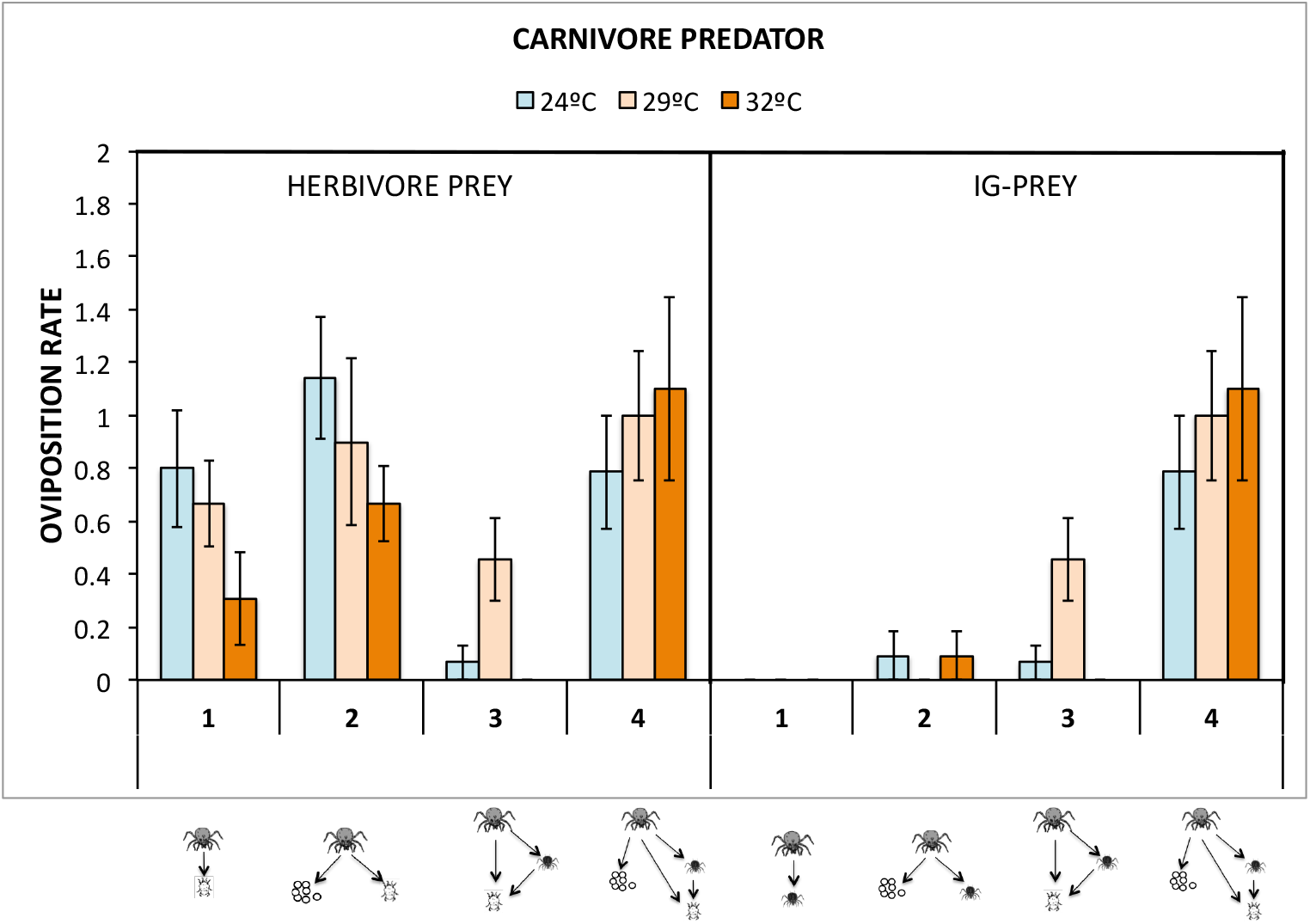
Oviposition rates of the *carnivore* predator when the focal prey was either the herbivore (left panel) or the *omnivore*-IGprey, measured in communities of increasing complexity (1 to 4 potential trophic links) and across increasing temperatures

### Predator-herbivore TIS: effects of temperature, presence of IG-prey, and presence of preferred food sources

When the predator was the *omnivore*, TIS exerted on the herbivore strongly depended on temperature and the presence of the *carnivore-IGprey* (Table 2a). Indeed, in the absence of IGprey the TIS exerted on the herbivore was very low and not affected by temperature (fig 2, left panel, compare communities 1 and 2 with 3 and 4, z= 3.299, P=0.005), whereas TIS strongly increased with temperature when the *carnivore-IGprey* was present (fig. 2, left panel, communities 3 and 4). This suggests that most of the trophic pressure on the herbivore was exerted by the juvenile *carnivore*-IG-prey, rather than by the *omnivore* top predator. Concomitantly, oviposition rates of the *omnivore* depended only on the presence of its preferred food source, pollen (Table 2b, figure 3, left panel, compare communities 1 and 3 with 2 and 4, t=7.861 P<<0,001). Yet, in the absence of pollen oviposition rates of the *omnivore* were still positive (fig 3 left panel, communities 1 and 3), indicating that predation on animal prey occurred.

**Table 2.**
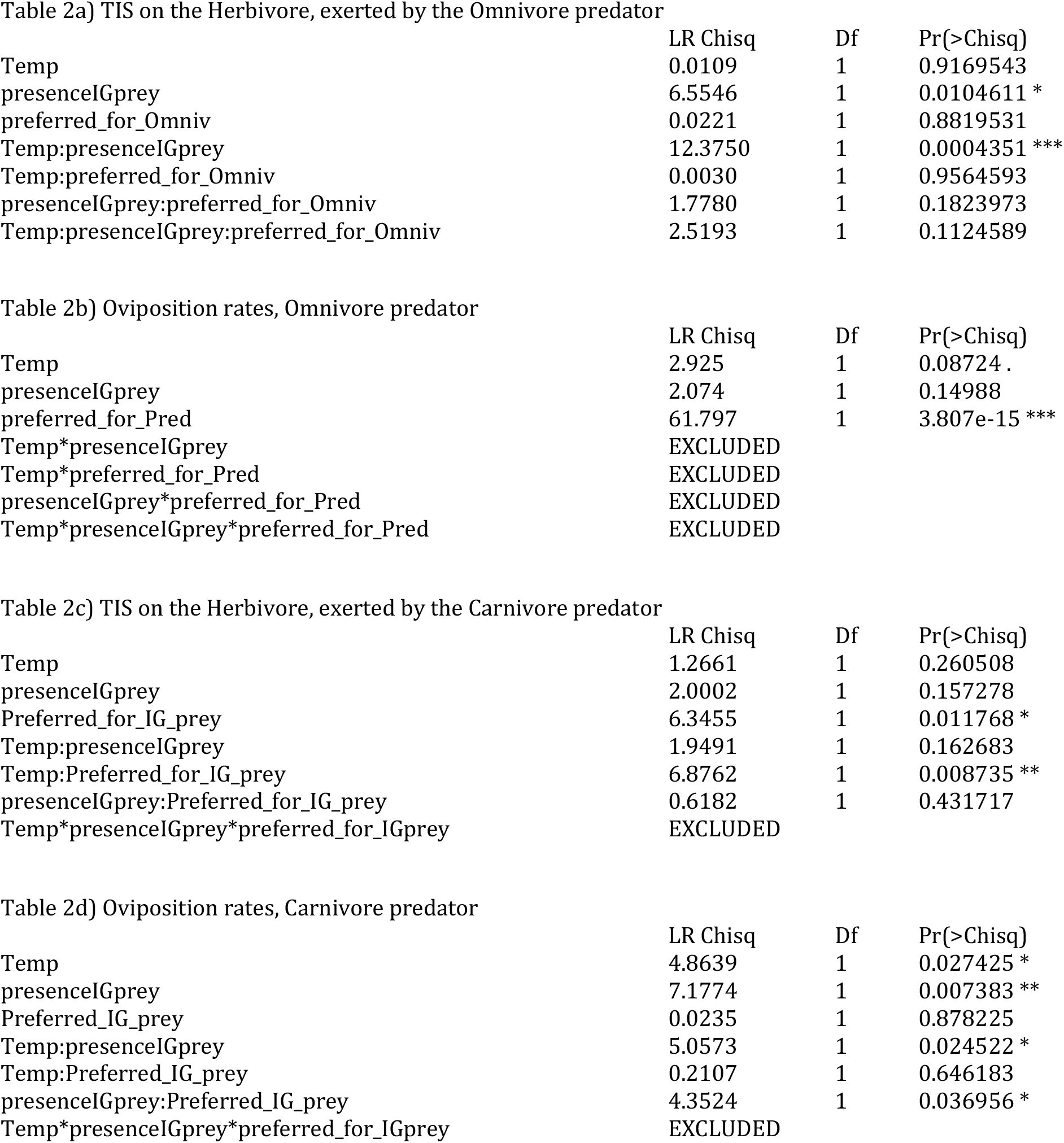
Results of the Generalized Linear Models applied to the dependent variables “trophic interaction strength” (TIS, with Gaussian distribution of errors) and predator’s “oviposition rates” (with Poisson distribution of errors). TIS on the <u>herbivore prey</u> was measured when they were exposed to either the omnivore (*E. stipulatus*, Table 2a and 2b, respectively) or the carnivore (*N. californicus*, Table 2c and 2d, respectively). In 2a and 2b, explanatory variables included in the models were “Temperature” as continuous variable, and “presence of IG-prey” (yes or not) and presence of the “preferred food source for the omnivore”, abbreviated as “preferred_for_Omniv” (yes or not), as categorical variables. In 2c and 2d, explanatory variables included in the models were “Temperature” as continuous variable, and “presence of IGprey” (yes or not) and presence of the preferred food source for omnivore IGprey, abbreviated as “preferred_for_IGprey” (yes or not), as categorical variables.

When the predator was the *carnivore*, TIS on the herbivore depended mainly on temperature and on the presence of the preferred food for the *omnivore*-IGprey (Table 2c). Indeed, in the absence of pollen and presence of the *omnivore*-IGprey TIS on the herbivore increased with temperature whereas no clear pattern was observed in the other communities (fig 4, left panel, compare community 3 with the others). Oviposition rates, however, depended on the interaction between temperature and presence of IGprey, and on the interaction between presence of IGprey and presence of preferred food for the IGprey (Table 2d). Indeed, in the absence of *omnivore*-IGprey oviposition rates of the *carnivore* decreased with increasing temperatures (fig. 5, left panel, communities 1 and 2), whereas in the presence of *omnivore*-IGprey there was no clear trend of temperature on oviposition rates (fig 5, left panel, communities 3 and 4). Also, in the presence of the preferred food for the *omnivore*-IGprey, oviposition rates of the carnivore were always higher than in its absence (fig 5, left panel compare communities 3 and 4, z=4.573, P<0.001).

### Predator-IGprey TIS: effects of temperature and presence of preferred food sources

When the predator was the *omnivore* TIS on the *carnivore-IGprey* depended on temperature and the presence of the preferred food source for the predator (Table 3a). Indeed, the trophic pressure of the *omnivore* on the *carnivore*-IGprey decreased with increasing temperatures, but only when the preferred food source for the *omnivore* was absent (fig. 2, right panel, compare communities 1 and 3 with 2 and 4). In the presence of pollen the *omnivore* mostly foraged on this food source (Table 3b and fig. 3, right panel, compare oviposition rates in communities 1 and 3 with those in communities 3 and 4, t=-4,286, P<<0,001).

**Table 3.** Results of the Generalized Linear Models applied to the dependent variables “trophic interaction strength” (TIS, Gaussian distribution of errors) on the IGprey, and predator’s “oviposition rates” (poisson distribution of errors). TIS on the <u>IG-prey</u> was measured when they were exposed to either the omnivore (*E. stipulatus*, Table 2a and 2b, respectively) or the carnivore (*N. californicus*, Table 2c and 2d, respectively). Explanatory variables included in the models were “Temperature”, as continuous variable, and “presence of the preferred food source for the predator”, abbreviated as “preferred_for_Pred” (yes or not) and “presence of the preferred food source for the IG-prey”, abbreviated as “Preferred_for_IGprey” (yes or not), as categorical variables.

|  | LR Chisq | Df | Pr(>Chisq) |
| --- | --- | --- | --- |
| Temp | 13.3399 | 1 | 0.0002598 *** |
| preferred_for_Pred | 18.3672 | 1 | 1.822e-05 *** |
| Preferred_for_IGprey | 0.0694 | 1 | 0.7921422 |
| Temp:preferred_for_Pred | 16.8343 | 1 | 4.079e-05 *** |
| Temp:Preferred_for_IGprey | 0.0969 | 1 | 0.7555957 |
| preferred_for_Pred:Preferred_for_IGprey | 0.0013 | 1 | 0.9711127 |
| Temp:preferred_for_Pred:Preferred_for_IGprey | 0.0004 | 1 | 0.9844514 |

Table 3b) Oviposition rate omnivore predator Response: Ovip
|  | LR Chisq | Df | Pr(>Chisq) |
| --- | --- | --- | --- |
| Temp | 1.058 | 1 | 0.3037 |
| preferred_for_Pred | 53.604 | 1 | 2.452e-13 *** |
| Preferred_for_IGprey | 0.036 | 1 | 0.8497 |
| Temp*preferred_for_Pred | EXCLUDED |  |  |
| Temp:Preferred_for_IGprey | EXCLUDED |  |  |
| preferred_for_Pred*Preferred_for_IGprey | EXCLUDED |  |  |
| Temp*preferred_for_Pred*Preferred_for_IGprey | EXCLUDED |  |  |

Table 3c) TIS on the omnivore IGprey, exerted by the Carnivore predator
|  | LR Chisq | Df | Pr(>Chisq) |
| --- | --- | --- | --- |
| Temp | 16.4582 | 1 | 4.973e-05 *** |
| preferred_for_Pred | 0.0264 | 1 | 0.8708785 |
| Preferred_for_IGprey | 8.0883 | 1 | 0.0044551 ** |
| Temp:preferred_for_Pred | 0.0049 | 1 | 0.9441453 |
| Temp:Preferred_for_IGprey | 13.3902 | 1 | 0.0002529 *** |
| preferred_for_Pred:Preferred_for_IGprey | 0.4434 | 1 | 0.5054743 |
| Temp:preferred_for_Pred:Preferred_for_IGprey | 0.7786 | 1 | 0.3775746 |

Table 3b) Oviposition rate Carnivore predator
|  | LR Chisq | Df | Pr(>Chisq) |
| --- | --- | --- | --- |
| Temp | 0.1819 | 1 | 0.6697 |
| preferred_for_Pred | 0.4098 | 1 | 0.5221 |
| Preferred_for_IGprey | 0.3661 | 1 | 0.5451 |
| Temp:preferred_for_Pred | 0.7724 | 1 | 0.3795 |
| Temp:Preferred_for_IGprey | 0.4856 | 1 | 0.4859 |
| preferred_for_Pred:Preferred_for_IGprey | 18.6377 | 1 | 1.581e-05 *** |
| Temp*preferred_for_Pred*Preferred_for_IGprey | EXCLUDED |  |  |

When the predator was the *carnivore*, results suggest that the predator basically did not engage in trophic interactions with the *omnivore*-IGprey. TIS ranged from zero to negative values with increasing temperatures (fig 5, right panel), indicating that mortality of *omnivore-IGprey* decreased when temperatures increased, especially in the presence of the preferred food source for the *omnivore*-IGprey (Table 3c, Fig 5, compare communities 1 and 2, z=2.767, P=0,029; and 3 and 4, z= 4.411, P<0,001). Oviposition rates indicate that the *carnivore* could forage and oviposit only in the presence of the herbivore and when the *omnivore*-IGprey’ preferred food source was available (Table 3d, fig. 5 right panel, compare community 4 with the others). This also suggest that *omnivore*-IGprey were likely interfering with the *carnivore* when they had no access to pollen.

## Discussion

In this work we explored the effects of increasing community complexity and temperature on predator: prey trophic interaction strength using an experimental system with two species of predatory mites and three potential prey types. Analyses revealed strong context dependent effects driven by predator and IGprey identity and by presence of preferred food sources for predators. Once they were accounted for, effects of temperature and community complexity on predator-prey TIS were unveiled. These can be summarized in three main points:

### Increasing community complexity does not necessarily reduce the strength of predator/prey trophic interactions

The effect of increasing community complexity on predator-herbivore TIS was clearly contingent on the identity of the predator. Indeed, when the top predator was the *omnivore* TIS on the herbivore was higher at the highest community complexities, but this was not the case when the top predator was the *carnivore* (Figure 2 and 4, left panels). In the case of the *carnivore*, this lack of effect confirms that the species *N. californicus* mostly forages on herbivore spider mites independently of the presence of other potential food sources (Pascua et al 2020). In the case of the *omnivore*, however, the result might appear counterintuitive as one would expect that the presence of different food sources would reduce the *omnivore*’s per capita predation rate on each of them. For example, the predatory mite *Amblyseius swirskii* consumed half of two herbivore prey [the whitefly *Trialeurodes vaporariorum* (Westwood) and the thrips *Frankliniella occidentalis* (Pergande)] when they were offered together than when each herbivore species was offered alone (Messelink et al 2008). This reduction in the per capita predation rate of predators in the presence of multiple food sources is the base of apparent mutualism (Holt and Lawton 1994), a positive indirect interaction between two prey species mediated by satiation of their shared predator, which is mostly observed during initial phases of population dynamics, before predators start responding numerically (Sydmonson et al 2002).

When the presence of preferred food sources was accounted for, results revealed that mortality of herbivores was mostly inflicted by the *carnivore-IGprey* (Fig. 2, left panel, communities 3 and 4), while the *omnivore*’s foraging and food conversion into eggs were contingent on the presence of its preferred food source, pollen, although some predation on animal food, and thus some conversion into eggs, also occurred in the absence of pollen (Fig 3, compare communities 1 and 3 with 2 and 4). These results suggests that when the *omnivore* and the *carnivore*-IGprey had access to their preferred food sources (pollen and spider mites, respectively) the structure of the community resembled one with two trophic chains, with each predator species foraging preferentially on its preferred food source, a result that matches that found in Torres-Campos et al. (2020). In the case of the *carnivore*, oviposition rates were, as expected, negligible in the absence of its preferred food source (spider mites) (Fig 5, right panel, communities 1 and 2), whereas in the presence of herbivore prey, oviposition rates of the *carnivore* were contingent on the presence of the preferred food source for the *omnivore-IGprey* (Fig 5, compare in both panels community 3 and 4). This suggest that *omnivore*-IGprey may have hampered the *carnivore’s* foraging on herbivores. Indeed, interference between predators promotes the persistence of primary consumers (Polis and Holt, 1992, Parshad et al 2016). Yet, interference vanished when *omnivore*-IGprey had access to its favourite food source, pollen. Again, this picture describes a situation where the community is reduced into two food chains (Torres-Campos et al., 2020).

These results suggest that the structure and persistence of communities might be strongly contingent on the presence of preferred food sources rather than on the number of species with potential to engage in predator-prey interactions. For example, theoretical works found that the parameter space for 3-species stable coexistence in systems with IG predation is enlarged when the omnivore switches prey following the diet rule (Krivan 2000, Krivan and Diehl 2005), that is, when inclusion of a food item in the diet depends on its relative profitability and abundance (Stephen and Krebs, 1986). Thus, partitioning of complex communities into simple communities with strong TIS driven by optimal foraging decisions could explain the persistence of some natural communities (Neutel et al. 2002, Montoya and Sole 2003).

### Warming strengthens predator-prey interactions and reduces conversion rates into eggs

Analyses accounting for potential sources of context dependence revealed that TIS of the *carnivore*-IGprey on the herbivore clearly increased with temperature (Fig 2, left panel, communities 3 and 4). This output has been extensively reported in the literature (e.g. Sentis et al 2014, Dell et al. 2013). While temperature had little effect on the TIS exerted by the *carnivore* on the herbivores (fig 4, left panel), oviposition rates of both the *carnivore* and the omnivore in most of the communities tended to decrease with warming (fig 3 and 5, left panels), supporting that in ectotherms the allocation of metabolic energy trades off with the components of the life history that determine fitness, that is, survival, growth, and reproduction (Burger et al 2019). Interestingly, this effect was mostly detected in those communities reigned by simple trophic chains.

### Presence of preferred food sources can counteract negative effects of warming on predator vulnerable stages

Mortality of IGprey was influenced by multiplicative effects among temperature, identity of predators, and community complexity. Indeed, when the predator was the *omnivore* TIS on the *carnivore*-IGprey was only existent at mild temperatures (Fig 2 right panel, communities 1 and 3 at 24°C). In the contrast, when the predator was the *carnivore* the survival of *omnivore*-IGprey increased with increasing temperature (Fig 4, right panel - note that TIS increases negatively). These results can again be better understood when the presence of preferred food sources is accounted for. On the one hand, the presence of pollen alleviated the mortality of *carnivore*-IGprey inflicted by the *omnivore* (fig 2, right panel, compare communities 1 and 3 with 2 and 4 at 24°C), supporting the hypothesis that *omnivores* focused on foraging on pollen. Also, in the presence of *omnivore*-IGprey, the *carnivore*’s TIS on the herbivore increased with warming only when the former did not have access pollen (Fig 4, left panel, compare communities 3 and 4). contradicting, at the two highest temperatures, the interference hypothesis exposed in the previous section. However, the performance of ectotherm predators usually worsens when temperatures exceed the optimal temperature at which foraging traits are maximal (Dell et al. 2011: Englund et al. 2011; Sentis et al. 2012, ref). Thus, in the absence of pollen, negative effects of temperature on the *omnivore*-IGprey could have alleviated interference, allowing the *carnivore* to resume foraging on spider mites (figure 4, left panel, community 3). However, a combined and stronger positive effect of temperature and presence of pollen on the *omnivore*-IGprey’s survival (fig 4, right panel, compare community 3 with 4) might have yielded some interference on the carnivore’s foraging, but low enough for it to be still able to prey on the herbivore (Fig 4, left panel, compare the decrease on TIS between communities 3 and 4 at the two highest temperatures). Thus, levels of interference between predator species may have depended on the presence of preferred food sources, but the magnitude of this indirect interaction also depended on temperature. Indeed, temperature-dependent interference was confirmed in behavioural observations that were done using this very same experimental system (Torres-Campos et al, in prep.).

### Consequences for the persistence of communities under climate change

Models based on metabolic theory show that warming affects the strength of trophic interactions (Rall et al. 2010; Sentis et al 2017), the length of food chains (Rall et al. 2010; Fussmann et al. 2014, Beveridge et al 2010), and the structure, size and stability of communities (Blanchard et al. 2012, O’Gorman et al. 2017, Uszko et al. 2017). Here, short term experiments were enough to detect, in our experimental system, that the presence of preferred food sources is key to better understand under what conditions warming may weaken or strengthen top-down control of a pest of avocado orchards. Firstly, in the absence of pollen our results indicate that at mild temperatures herbivore top-down control might be hampered via both predation of the *omnivore* on the *carnivore-IGprey*, and interference between the *omnivore*-IGprey and the *carnivore*. Warming, however, could increase herbivore control by the *carnivore*, reducing both predation and interference by the *omnivore* via negative effects of temperature on the performance of both *E. stipulatus* adults and juveniles (Guzman et al 2018). Thus, in the absence of pollen, warming could strengthen top-down control of the herbivore mite pest. Other experimental works, however, found that warming benefits intraguild predator populations by increased predation rates on consumers (Frances and McCauley 2018; Rogers et al. 2018). Secondly, in the presence of pollen our results indicate that at mild temperatures resource partitioning in the two predator species, leading to a community with two food chains, could strengthen the herbivore top-down control by one of them. With increasing temperatures, however, pollen increasing the survival and the activity levels of the *omnivore*-IGprey could increase intraspecific interference and reduce the efficiency of the *carnivore*. Thus, in the presence of pollen, warming could weaken top-down control of the herbivore pest.

Our data supports that the effects of warming on food web structure may largely depend on how prey preferences of predators affect species interactions within communities (Lindmark et al 2019), and that non consumptive effects of predators on prey, and viceversa, can interact with warming and be as important as direct feeding (Preisser et al. 2005, Janssens et al 2015, Ecology). Thus, we conclude that that sources of context dependence inherent to the identity and characteristics of the species that compose the communities under study, need to be identified (Catford et al 2022) to unveil obscuring effects of warming because interactive responses to temperature and to other components of the community might be species-specific (Walker et al 2020). Thus, studying the diet breadth and food preferences of the species embedded in communities may be more important than previously thought to understanding the role of temperature and community complexity on the overall responses of populations, communities and associated ecosystem processes to global warming.

